# Functional Interpretation of Single-Cell Similarity Maps

**DOI:** 10.1101/403055

**Authors:** David DeTomaso, Matthew Jones, Meena Subramaniam, Tal Ashuach, Chun J. Ye, Nir Yosef

**Affiliations:** Center for Computational Biology, University of California Berkeley, Berkeley, CA, USA; Biological and Medical Informatics Graduate Program, University of California, San Francisco, CA, USA; Department of Epidemiology and Biostatistics, Department of Bioengineering and Therapeutic Sciences, Institute for Human Genetics, University of California, San Francisco, CA, USA; Department of Electrical Engineering and Computer Science and Center for Computational Biology, University of California, Berkeley, Berkeley, CA, USA; Ragon Institute of Massachusetts General Hospital, MIT and Harvard, Cambridge, MA, USA; Chan Zuckerberg Biohub Investigator

## Abstract

We present VISION, a tool for annotating the sources of variation in single cell RNA-seq data in an automated, unbiased and scalable manner. VISION operates directly on the manifold of cell-cell similarity and employs a flexible annotation approach that can operate either with or without preconceived stratification of the cells into groups or along a continuum. We demonstrate the utility of VISION using a relatively homogeneous set of B cells from a cohort of lupus patients and healthy controls and show that it can derive important sources of cellular variation and link them to clinical phenotypes in a stratification free manner. VISION produces an interactive, low latency and feature rich web-based report that can be easily shared amongst researchers.

## Background

Recent technological advancements have allowed transcriptional profiling at the single-cell level [1, 2, 3]. This has enabled a deeper investigation into cellular heterogeneity [4], the identification of new cellular subtypes [5, 6], and more detailed modeling of developmental processes [7, 8]. Notably, the data produced in a single-cell RNA-seq (scRNA-seq) experiment is distinct from that of bulk RNA-seq in that it is typically sparse (with many expressed genes remaining undetected), and consists of a very high number of data points [9]. Furthermore, while bulk studies are usually conducted in a comparative setting (e.g., case control studies), most investigations of scRNA-seq are based on comparisons of cells within a sample, without any preconceived labels of these cells.

A typical primary step in the analysis of scRNA-seq data is therefore to extract a meaningful labeling by partitioning the cells into groups [10, 11] or by placing the cells along some continuum [12] in a data-driven manner. A common way to achieve this is to first project the data onto a low-dimensional space, which preserves critical information while reducing noise and bias. While Principal Component Analysis (PCA) is a commonly used projection method, more recently linear factor models such as ZIFA[13] or zinbWave[14] and nonlinear deep generative models such as scVI[11] or DCA[15] have been developed to specifically address the underlying distributions and confounders found in single-cell RNA-sequencing. The resulting manifold representations [16] can then be used as a baseline for dividing the cells into clusters. Alternatively, if the cells are expected to vary along a continuum, such as that which arises during a developmental time-course, a tree-like representation of the data can be inferred instead (summarized in [12]).

While the assignment of labels (e.g., cluster IDs) to cells greatly simplifies the interpretation of the data, it may come at the cost of missing important yet subtle patterns of variation (e.g., gradients of important cellular functions within a cluster of cells [17]) and suffer from inaccuracies (e.g., when there is no obvious cluster structure [18]). Furthermore, even once labels have been assigned, it may still not be clear how to interpret their underlying biological meaning. To address these challenges and identify the key biological properties that dominate the variability between cells in a sample, we developed VISION, a flexible annotation tool that can operate both with and without preconceived labeling of cells (Figure 1a). As an input VISION takes a map of similarities between cells, which can be computed internally or provided by external manifold learning algorithms [16, 13, 14, 11]. VISION then leverages the concept of transcriptional signatures [19, 17] to interpret the meaning of the variability captured in the manifold by integrating information from a large cohort of published genome-scale mRNA profiling datasets [20, 21, 22]. In its label-free mode, VISION operates directly on the single cell manifold and uses an autocorrelation statistic to identify biological properties that distinguish between different areas of the manifold. The result is a set of labellings of the cells, which may differ when studying different aspects of cell state (e.g., tissue context vs. differentiation stage in T cells [18]). This approach is therefore capable of highlighting numerous gradients or sub-clusters which reflect different cellular functions or states and which may not be captured by a rigid precomputed labeling. As we demonstrate, this approach is particularly helpful when studying cells from the same type (e.g., naïve B cells), with no clear partitioning. In its label-based mode, VISION identifies biological properties that differ between precomputed stratifications (e.g., clusters) or that change smoothly along a given cellular trajectory. To enable the latter, VISION utilizes the API built by Saelens and colleagues [12] to support a large number of trajectory inference methods, and it is the first functional-annotation tool to do so.

**Figure 1:**
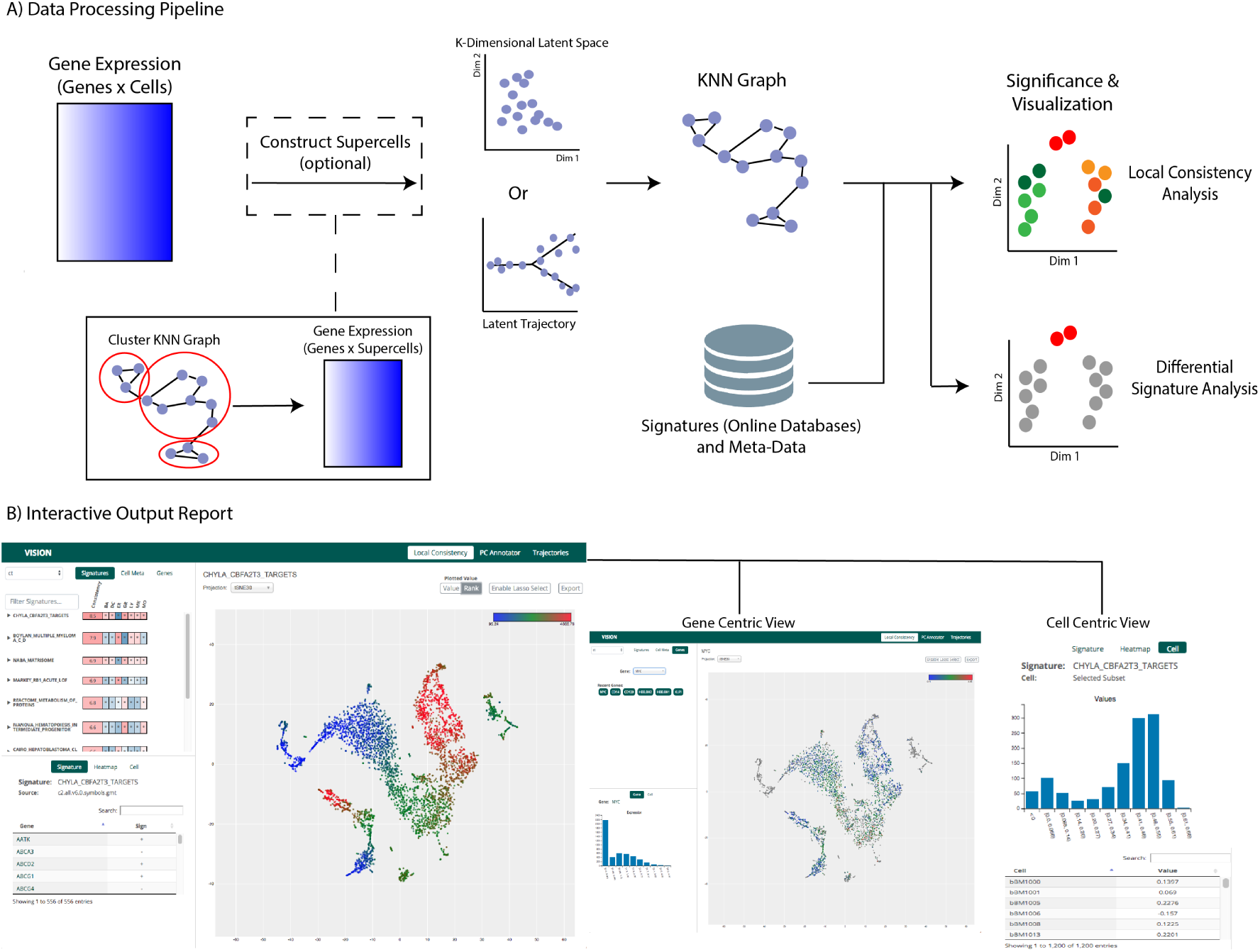
A) The VISION processing pipeline consists of several key steps. A K-nearest neighbors graph is constructed within the latent model for gene expression (either supplied as input, or computed using PCA). Optionally, this graph can be reduced for more efficient downstream computation by combining similar cells into ‘micro-clusters’. Representative scores are evaluated for external gene signatures and local autocorrelation analysis is used to evaluate which signatures best reflect the heterogeneity in the latent space. B) The analysis output is organized into an interactive report in which signatures, gene expression, and cell meta-data can be visualized along with two-dimensional representations of the data.

VISION has several additional properties that distinguish it from other software packages for automated annotation and for visualization and exploration of single cell-data (summarized in Supp. Table 1). Foremost, VISION is designed to naturally operate inside of analysis pipelines, where it fits downstream of any method for manifold learning, clustering, or trajectory inference and provides functional interpretation of their output. VISION also enables the exploration of the transcriptional effects of meta-data, including cell-level (e.g., technical quality or protein abundance [23]) and sample-level (e.g., donor characteristics). As we demonstrate, VISION identifies the degree to which library quality drives cellular heterogeneity as well as illuminates the relationship between clinical quantities of inflammation burden and transcriptional properties of immune cells. Finally, the use of VISION can greatly facilitate collaborative projects, as it offers a low-latency report that allows the end-user to visualize and explore the data interactively. The report can be hosted on-line and viewed on any web browser without the need for installing specialized software (Figure 1b). VISION is freely available as an R package at www.github.com/YosefLab/VISION.

## Using Signature Scores to Interpret Neighborhood Graphs

VISION operates on a low dimensional representation of the transcriptional data and starts by identifying, for each cell, its closest K-nearest neighbors in the respective manifold. Computing this for every cell results in a K-nearest-neighbor (KNN) graph. By default, VISION performs PCA to create this low dimensional space, but the results of more advanced latent space models [13, 14, 11] or trajectory models (via [12]) can be provided as an input instead (to note, these trajectory models may be described as both latent spaces and a precomputed labeling of the cells). In order to interpret the variation captured by the KNN graph, VISION makes use of gene signatures - namely, manually-annotated sets of genes, which describe known biological processes [24] or data-driven sets of genes that capture genome-wide transcriptional differences between conditions of interest [25]. These signatures are available through databases such as MSigDB [26], CREEDS [21], or DSigDB [22] and can also be assembled in a project-specific manner (e.g., as in [17, 27]). For each signature, an overall score is computed for every cell summarizing the expression of genes in the signature. For example, for a signature describing inflammatory response, a high signature score would indicate that the cell in general has higher expression of known inflammatory response genes. Gene signatures may also be ‘signed’ - representing a contrast between two biological conditions. For example, given a signature representing Th17 helper cells vs. regulatory T cells, a higher score would indicate that a cell’s transcriptional program is more Th17-like, while a lower score would indicate it is more similar to the regulatory state [17, 28]. To reduce the effect of batch or technical covariates on these signature scores, we recommend the use of a normalization procedure (such as in [29], [11] or [30]) on the gene expression dataset prior to input into VISION.

To interpret the cell-cell KNN graph in the context of signature scores without the use of labels (namely, label-free mode), we make use of a local autocorrelation statistic, the Geary’s C [31]. This statistic was originally developed for use in demographic analysis to identify significant spatial associations (e.g. “Are incident rates of obesity randomly distributed within a city or is there a certain areal pattern?”). Here, we make use of this statistic to answer a similar question: “Is the signature score randomly distributed within the KNN graph or is there a pattern?” (see Methods for a complete description of this statistical test). Signatures with a significantly high local autocorrelation statistic can therefore be used to assign a biological meaning to specific areas of the KNN graph, and also capture gradients or various sub-divisions along the graph. This is especially useful for cases where the cells do not clearly separate into discrete clusters, but rather exhibit variation in a more continuous fashion. Since oftentimes significant biological factors correlate with principal components [17, 32], VISION offers an additional label-free analysis, dubbed “PCAnnotator”, which allows the user to rank signatures according to correlations with the individual components of the latent space. In its label-based mode VISION evaluates the dependence of the biological signatures on the labels assigned to each cell, such as experimental group or cluster ID. To accomplish this, VISION uses a 1-vs-all differential signature test (see Supplementary Methods) to highlight biological properties that distinguish each cluster.

In addition to gene signatures, VISION allows the user to directly input other properties (or meta-data) for each cell, and explore their effects in a similar manner to that of the gene signatures. This approach can be useful for evaluating the degree to which quality control parameters (e.g., % of mapped reads) are driving cellular heterogeneity. It also allows the integration of additional sources of biological information such as protein abundance per cell [23] or sample-level attributes such as time point or donor clinical parameters. The meta-data may also be categorical and represent properties such as batch annotations or specifications of the respective experimental condition. VISION enables the analysis of these categorical properties in both local autocorrelation (label-free) and comparative (label-based) modes using a chi-squared test.

## Distinct Signature Modules Drive Transcriptional Variation in B Cells of Lupus Patients

As a demonstration, we applied VISION to a new single-cell RNA-seq dataset consisting of peripheral blood mononuclear cells (PBMCs) derived from a cohort of twelve Systemic Lupus Erythematosus (SLE) patients and four healthy controls (see Methods for description of samples and protocols). Initially, we utilized VISION to analyze the complete collection of 67k PBMCs. Because the cells exhibit clear, cell-type differences, we used the unsupervised clustering performed by VISION to partition the cells into labeled groups. By employing a set of cell-type signatures and performing label-based, 1 vs. all differential signature tests, these labeled groups can be contextualized into broad annotations (e.g., CD4+, CD8+ T cells, monocytes etc. Supplementary Figure 2a,b).

Additionally, further variation exists *within* cell-type clusters and the label-free signature analysis performed by VISION can assign a functional interpretation to this heterogeneity. To demonstrate this, we subset the data to focus on the B cell cluster (6,857 cells). The signatures found to have significant local autocorrelation within the B cells (false discovery rate (FDR) <0.05) are automatically partitioned in the VISION output report into distinct groups based on the similarity of their respective scores (namely, similar signatures capture similar gradients in the data). One module is characterized by signatures that describe the memory vs naive contrast (Figure 2c) revealing that the B cell cluster can be further subdivided according to these types. Notably, the differences between these two sub-types is more subtle in comparison to the differences between cell types and may therefore not be easily captured in an unsupervised pan-PBMCs clustering.

**Figure 2:**
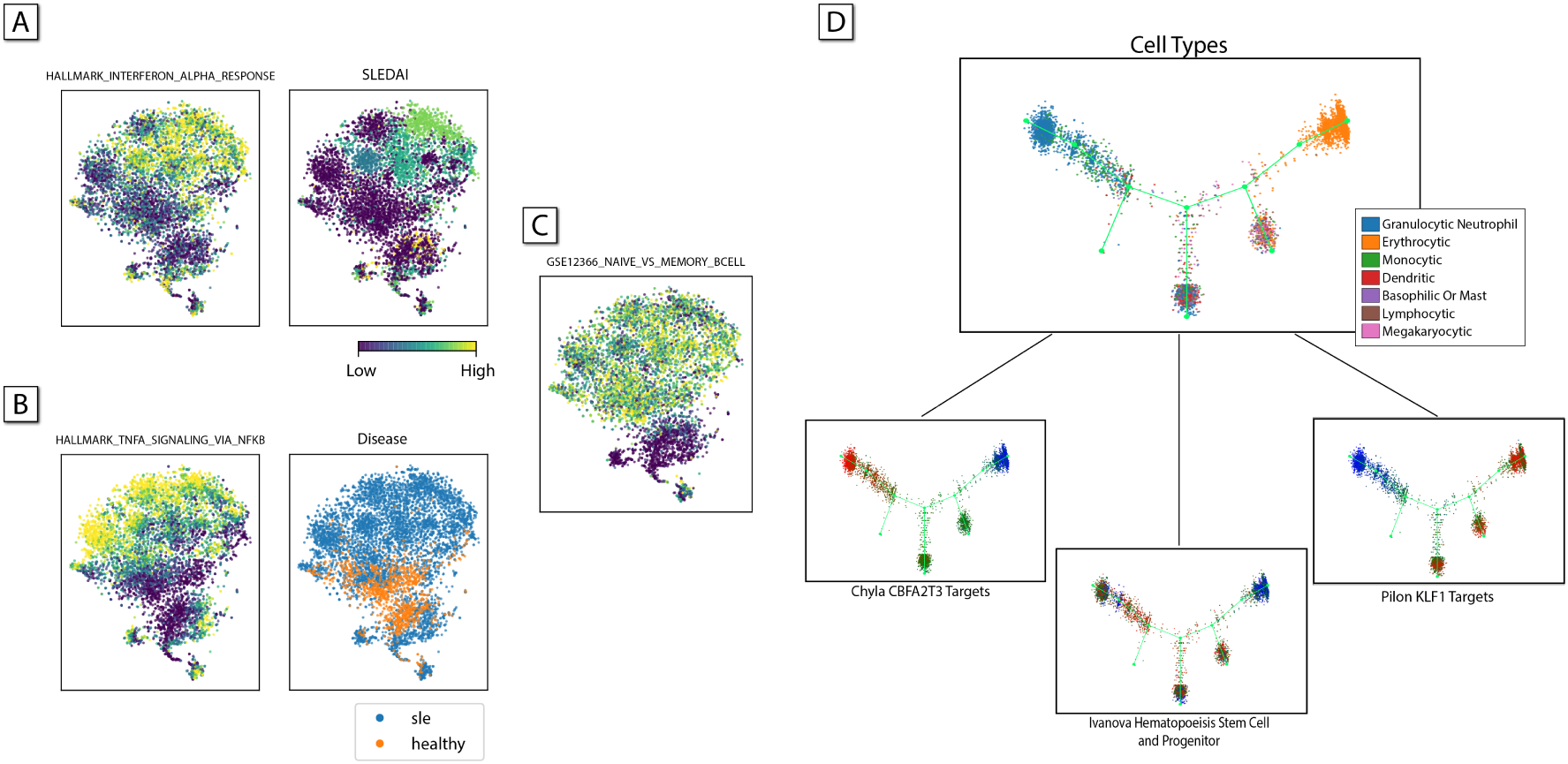
A) VISION identifies signatures describing transcriptional variation within B cells in a mixed cohort of healthy and SLE patients. Clustering the signatures with high local autocorrelation reveals distinct modules of variation which can be characterized by memory (C), Interferon Alpha (A), and Tumor Necrosis Factor Alpha (B) signatures. In (A), the SLEDAI score is observed to exhibit a similar pattern of variation to signatures in the Interferon Alpha cluster while in (B) the Disease status is highly indicative of a patient’s cell’s TNF*α* signature score. D) Signatures are used to describe variation within the context of a trajectory model for haematopoietic differentiation. High-scoring transcriptional signatures reflect help to annotate trajectory branches and highlight transcription factors involved in lineage commitment.

Two other signature groups highlight different, intriguing patterns of variation that are present in both sub-types of B cells. The first group consists of signatures that are indicative of pro-inflammatory activity, such as genes associated with Interferon-*α* Response (Hallmark collection, MSigDB[26]), a known key cytokine in the pathogenesis of SLE[33]. The second group includes a signature of TNF*α* response genes (Hallmark collection, MSigDB[26]), which is known to increase in patients with SLE [34]. To better understand what these distinct patterns of variation mean, we sought to contextualize their scores by adding donor-level meta-data. Using disease status (case/control), the VISION label-free mode finds, as expected, that the similarity between cells strongly depends on the disease status of the respective donor (Chisq test, *V* = 0.41, *p* < 10^−20^).Using the label-based mode, namely searching for signatures that significantly differ between SLE and controls, VISION reports that both TNF*α* and Interferon-*α* are strong indicators for disease status (adjusted Wilcoxon rank-sum *p* < 10^−20^). However, closer inspection reveals that TNF*α* is a stronger indicator (AUC 0.81 vs 0.74), while high expression of Interferon-*α* genes characterizes only a subset of the SLE-derived B cells (Figure 2b).

As a second source of meta-data, we used the SLEDAI score - a quantitative aggregate clinical measure of disease symptoms. As before, using the label-free mode, VISION finds this quantitative score to be a strong indicator of cell-cell similarity (autocorrelation *C′* = 0.45, *p* < 0.0038). Since this is a quantitative score, VISION also automatically clusters it with the other, data-driven, gene signatures. In this case, the SLEDAI score is assigned to the same group as Interferon-*α* based on cell-level correlation. (Figure 2a). Closer inspection of the scores reveals that Interferon-*α* serves as a better predictor of SLEDAI score compared to TNF*α* even after considering only SLE donors and controlling for cell to donor association (*p* < 0.048 vs. *p* < 0.82, see supplementary methods). Overall, this suggests two independent modes of transcriptional variation: one which better distinguishes healthy and sick donors (characterized by TNF*α*) and the other which more closely corresponds with the disease burden (characterized by Interferon-*α*).

To further probe the effects of disease status on the B cell population, we used the label-based approach to investigate transcriptional signatures associated with cytokine signaling (NetPath database [35]). We find that the IL-3 and Wnt signatures emerge as the most discriminating between SLE and control donors (Supplementary Figure 2e). Notably, previous studies support a role for IL-3 in SLE in connection with B cells: IL-3 is a growth factor for B cells[36], serum levels of IL-3 have been observed to be increased in SLE patients vs healthy controls[37], and a recent study in a lupus nephritis mouse model found that IL-3 blockade ameliorated disease symptoms and notably reduced splenic B-cell counts[38].

VISION can be additionally used to assess the degree to which other forms of sample and cell-level meta-data is reflected in the underlying cell-cell similarity map. Applied to this data, it can be observed that within the B cells, the local distribution of the donor covariate is highly significant (*V* = 0.34, *p* < 10^−20^ Supplementary Figure 2c), demonstrating that the donor-donor heterogeneity is more significant than the cell-cell heterogeneity in this data. Alternately, it can be seen that the 10x well covariate is well mixed in the similarity (*V* = 0.01, *p* < .99 Supplementary Figure 2d).

Finally, VISION’s PCAnnotator feature highlights much of the same underlying biology discussed here. Preliminarily, we observed that the first PC stratifies well the B cells from cases and controls, indicating that the principal components might be capturing biology driving this stratification (Supplementary Figure 4a). The results from the PCAnnotator support this hypothesis, as we find that the first PC correlates with both signatures describing the TNF*α* response as well as the Interferon-*α* response (*ρ* = 0.73 and *ρ* = 0.46 respectively, Supplementary Figure 4b). Overall, the combination of biological signatures, pre-computed labels, meta-data, and principal components can be used to distinguish some of the key biological features driving heterogeneity in this collection of B cells.

## Annotating Cellular Trajectories: Haematopoietic Differentiation as a Case Study

Our scheme for biological interpretation with local autocorrelation can also be applied to cell-labellings from trajectory maps, asking: “Is there an association between the position of a cell in the trajectory and a certain biological function?” To facilitate this test, VISION computes a KNN graph from pre-computed trajectory models, connecting cells that are close to each other in the inferred continuum. The autocorrelation statistic is then computed on the KNN graph in a manner similar to the analysis above. Importantly, VISION supports a variety of trajectory inference methods through integration with the Dynverse[12] package, which provides wrappers for approximately thirty different algorithms.

To demonstrate the utility of this module, we used Monocle2 [7] on 5432 haematopoietic progenitor cells (HPCs) isolated from the bone marrow of adult mice [39]. VISION’s rendering of the Monocle output recapitulates the pattern of differentiation and the stratification of cells discussed in the original study - namely, an undifferentiated core giving rise to differentiated cells, most notably erythrocytes and granulocytes (Supplementary Figure 3). We used the Hallmark (H) and curated gene set (C2) collections from MSigDB [26] to attribute additional meaning to the differentiation process. Unsurprisingly, signatures distinguishing granulocytes and erythrocytes (Neutrophil Granule Constituents (*C′* = 0.7, FDR < 4.5 × 10^−3^) and Hallmark Heme Metabolism (*C′* = 0.72, FDR < 4.5 × 10^−3^), respectively) highlighted the granular neutrophil and erythrocytic arms of the trajectory. Furthermore, high values for a signature describing haematopoietic stem cell and progenitor populations (Hematopoiesis Stem Cell and Progenitor, *C′* = 0.4, FDR < 0.04) significantly localize to the undifferentiated core of the trajectory (Figure 2D).

Notably, additional interesting signatures showed to be significant, emphasizing more nuanced biological processes occurring during hematopoiesis. For example, the granulocytic arm showed high signature scores for the CBFA2T3 Targets signature (*C′* = .89, FDR < 4.5 × 10^−3^). This signature includes genes that are up-regulated after Mgt16 knockdown, skewing HPCs to granulocytic/macrophage lineages and thus highlighting Mgt16 as a key regulator of HPC lineage commitment. Additionally, the KLF1 Targets (*C′* = 0.83, FDR < 4.5 × 10^−3^) signature, which includes genes that are potential EKLF targets responsible for the failure of erythropoiesis, illustrates clearly that KLF1 is an important regulator of the erythrocytic lineage. Taken together, these results show that the combination of signatures and latent trajectory models can emphasize important regulators of dynamic processes such as development.

As a complementary approach, VISION can perform analysis on numerical meta-data, such as quality control (QC) measures. We make use of this function to identify numerical signatures which best distinguish different arms of the haematopoietic trajectory. Firstly, we observed that the ratio of genes detected in a cell (referred to as the cell detection ratio, or CDR) and the number of UMIs were overall locally consistent as determined by the Geary C statistic (*C′* = 0.93 and *C′* = 0.90 respectively, FDR < 4.5 × 10^−3^ for both). Then, leveraging the label-based differential signature test, we find that these two meta-data signatures are both associated with higher values in granulocytic cells (FDR < 4.5×10^−3^, both, Supp. Figure 3b). This likely reflects that granulocytes tend to have diameters twice the size of other white blood cells and erythrocytes. However, in other experiments where such a difference is not expected, such a result may indicate the data is confounded by technical noise and requires further correction through various forms of scaling or normalization[29, 14].

## Discussion

Here we have presented VISION - a tool for the functional interpretation of cell-to-cell similarity maps. VISION builds upon our previous work, FastProject [19], which was designed to annotate two dimensional representations of single cell data. Overall, the work described here provides a substantial increase in functionality such as support for higher-dimensional latent spaces, explicit interpretation of pre-computed clusters, cellular trajectories, and other meta-data, as well as improved scalability. VISION also refines FastProject’s core algorithms, most notably including a new signature autocorrelation score statistic (Methods). Because VISION is able to operate on a variety of manifolds (latent spaces or trajectories from a broad array of methods) and scale to a large number of cells, it is well suited to benefit the interpretation of single-cell RNA-seq data as modeling methods continue to evolve. Moreover, VISION offers a more unified approach and greater flexibility to researchers because it may sit downstream of other methods, thus decoupling the choice of modeling algorithms and interpretation.

The results of a VISION analysis can be explored through the use of an interactive web-based GUI (Figure 1b). In this interface, top-scoring signatures are listed and can be visualized on the single cell manifold through the use of two-dimensional projections such as those generated with t-distributed Stochastic Neighborhood Embedding [40]). If the latent space is a trajectory instead, the tree structure is visualized along with cells using various algorithms (e.g. Davidson-Harel, Fruchterman-Reingold) for tree embedding in two dimensions. To improve the interpretability of the expression data, VISION’s output report offers four possible views: a “signature-centric” view which highlights modulation of between groups of the data; a “cell-centric” view which allows users to independently analyze a subset of cells; a “gene-centric” view which allows users to view a single gene at a time; and the PCAnnotator which dissects the relationship between principal components and biological heterogeneity. These features can help identify either important cell subtypes, crucial genes, or biological factors driving variation seen at the population level.

VISION can accommodate 50,000 cells in around 30 minutes (Supplementary Figure 1a); however, to scale well into the hundreds of thousands of cells, VISION utilizes a micro-pooling algorithm in which the expression profiles of nearby cells in the similarity map are merged into representative “micro-clusters” (see Methods). Overall, we find that the local autocorrelation of signatures remains consistent after micro-pooling and that the micro-clusters produced are biologically coherent (Supplementary Figure 1b, c). Taken together, we find this approach heavily reduces the computation time - allowing the analysis of 500,000 cells with 20 cells per micro-cluster in under an hour while producing results consistent with the non-pooled analysis.

In comparison with other single-cell RNA-seq software tools, VISION augments the functionality of toolkits like Seurat [41] and Scanpy [42], which can be used for normalization and labeling of the gene-expression data prior to VISION analysis. VISION is distinct from visualization tools such as SPRING [43] in that it offers functional interpretation of the single-cell data. Moreover, VISION goes beyond standard workflows that provide gene set enrichment analysis on genes differentially-expressed between groups (such as that offered by MAST [44]), by providing functional interpretation for inferred trajectories and for cases when the cells cannot clearly be clustered into groups. Finally, in comparison to tools that can annotate important axes of biological variation without the need for a priori stratification such as PAGODA ([45]) and ROMA ([46]), we illustrate that VISION’s signature scores more effectively capture the underlying biological differences between samples (Supp. Figure 5) and more precisely highlight crucial variation and sub-clusters in the data (Supp. Figure 6, see methods). A comparison of the analysis features available in VISION versus those available in other tools is provided in Supp. Table 1.

In summary, VISION offers a high-throughput and unbiased method for profiling variation and heterogeneity in single cell RNA-seq data. As the number of methods for both processing and obtaining single cell measurements (e.g. CITE-seq for simultaneous protein and gene measurements [23]) increases, we anticipate that methods like VISION which are able to integrate different data types and flexibly interpret the results of any analytical method will be in high demand. Finally, because the results of VISION can be made available through an interactive web-based report, we expect that VISION will allow for improved reproducibility and communication of scRNA-seq results. Given these features, we anticipate VISION’s widespread usage as a tool for functional interpretation, technology integration, and collaboration.

## Supplementary Methods

### Signature Score Calculation

A signature is defined as a set of genes associated with some biological function or measured perturbation. Signatures may be *signed* in which there are two sets of genes, a positive set *G_pos_* and a negative set *G_neg_*. Such a signature is used when describing an experimental perturbation or a comparison between two cell states in which some genes increase in expression while others decrease. Alternately, a signature may be *unsigned* in which case *G_neg_* is empty.

For each signature, a representative score is computed for every cell. This is calculated as the sum of expression values for positive genes minus the sum of expression values for the negative genes. For example, for signature *s* and cell *j* the score is computed as:

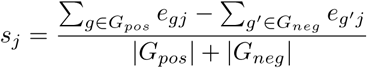

In the expression measure above, *e_gj_* is taken to be the normalized, log-scaled expression (e.g. log of counts-per-million + 1 or log of counts-per-thousand + 1) of gene *g* in cell *j*. However, we have observed that even after performing standard normalization procedures on the expression values (e.g., regressing out technical covariates), signatures scores as defined above still tend to be correlated with sample-level metrics (such as the number of UMIs per cell). To account for this, we z-normalize the signature scores using the expected mean and variance of a *random* signature with the same number of positive/negative genes. Specifically, for a signature score, *R_j_* in cell *j* derived from a random signature with *n* positive genes and *m* negative genes:

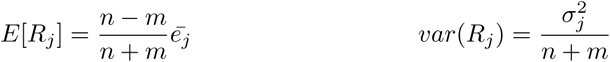

where 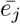 and 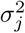 represent the mean and variance respectively of the expression values for cell *j*.

The final, corrected signature score, 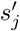 is computed as:

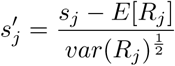

### Local Autocorrelation Calculation

To compute the extent to which a signature can explain the variation in a cell-to-cell similarity map, we make use of the Geary’s C statistic for local autocorrelation. This statistic is defined as:

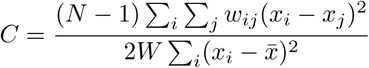

where *w_ij_* represents the weight between samples *i* and *j* in some map, *x_i_* is a value of interest, *N* is the total number of samples, and *W* is the sum of all weights. In our case, the value of interest (*x* values) are the ranks of the normalized signature score in each sample. The weights, *w_ij_* between samples (cells) *i* and *j* are set to be nonzero and positive for cells nearby in the provided latent space (details follow).

In this way, the Geary’s C provides a measure of how similar the signature ranks are for neighboring cells given a latent mapping. For the interactive output report, we report *C′* = 1 *− C* as the autocorrelation effect size so that a 0 intuitively represents no autocorrelation and a 1 represents maximal autocorrelation.

To compute the cell-cell weights, *w_ij_*, for the Geary’s C, first the *k* = *N* ½ nearest neighbors are evaluated for each cell in the provided latent map. If the input map is a latent space, this is evaluated using euclidean distance in the latent space. If the input map is a tree or trajectory, this is evaluated using the path distance along the trajectory. To accommodate the variety of latent trajectory methods which have been developed for single-cell RNA-seq, we make use of the Dynverse package[12] which rectifies the output of 29 such methods to a common trajectory model.

Once distances and neighbors have been determined, weights can be calculated. For cells which are k-nearest neighbors, the weight is evaluated as:

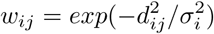

where *d_ij_* is the distance between cell *i* and *j* in the latent mapping and 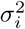 is the distance to the kth-nearest neighbor of cell *i*. If cell *j* is not a k-nearest neighbor of cell *i*, then *w_ij_* is taken to be 0.

### Assessing Significance of Autocorrelation Scores

To evaluate the significance of the autocorrelation scores for each signature, a set of random signatures are generated (genes drawn from the set of genes in the input expression matrix), and autocorrelation scores on these signatures are computed to act as an empirical background distribution. The p-value for signatures is then computed as 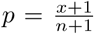 where *x* is the number of background signatures with a higher autocorrelation score and *n* is the total number of background signatures. These p-values are then corrected for multiple-testing using the Benjamini-Hochberg procedure.

To avoid the computational cost of generating a random background for every evaluated signature, we instead create 5 background signature groups which encompass the range of signature sizes (number of genes) and balance (ratio of pos/neg genes) in the input signature set. This is suffcient as we have observed that the distributions of random signature p-values are very similar even when the size and background are not perfectly matched. The size and balance of background signature groups is evaluated by clustering all input signatures by their *log*_10_(*size*) and *balance* using k-means with *k* = 5. Cluster centers are then used for the background group sizes and balance, and cluster assignments are used to match signatures under test with the closest background.

### Micro-pooling

VISION employs a micro-pooling algorithm to partition the input expression matrix and pool cells together, resulting in a reduction of computational burden for data sets consisting of a large number of cells. The algorithm begins by applying gene filters to the input expression matrix: genes are first thrown out that are not expressed in at least 10% of cells and then highly variable genes are selected as in [19]. Next, we project the filtered matrix down to 20 dimensions using PCA. Then for the *N* cells in the expression matrix, the K-nearest-neighbor graph (with *K* = *sqrt*(*N*)) is computed in this reduced space.

Initially, this KNN graph is clustered using the Louvain algorithm, an effcient community detection algorithm. These clusters are further partitioned with K-means until each cluster consists of at most *P* cells per partition (*P* can be controlled by the user).

“Micro-clusters” are then generated using these partitions. For each partition, we create a micro-cluster whose gene expression values are defined as the average gene expression values for each cell in the partition. More specifically, for gene *i* in micro-cluster *z* generated from partition *P_z_*, the expression value for this gene *e_iz_* is equal to:

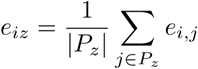

Finally, an expression matrix consisting of these micro-clusters of dimension *O*(*N/P*)*×G* is returned and used for downstream analysis.

### Assessing Biological Coherence for Micro-Clusters

We assessed the biological coherence of the micro-clusters with a dataset consisting of simultaneous epitope and gene expression profiles of single cells, published in [23] (Gene Expression Omnibus accession GSE100866). For ~ 9, 000 Cord Blood Mononuclear Cells (CBMCs), we performed micro-pooling on the *transcriptional* data to create micro-clusters with at most 20 cells per cluster with the gene expression data. Then we analyzed the relative variation in *protein* within each micro-cluster. More specifically, we reported the ratio of intra-micro-cluster standard deviations to the overall standard deviation of the protein across the entire population of cells:

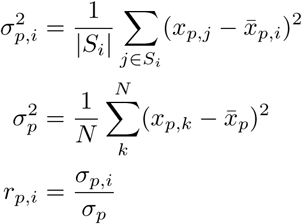

where 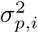 is the variance across the cells in micro-cluster *i* for protein *p*, 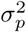 is the population-wide variance of protein *p, x_p,j_* is the abundance of protein *p* in cell *j*, 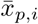 is the mean abundance of protein *p* across cells in micro-cluster *i*, 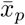 is the mean abundance of protein *p* across the population, and *r_p,i_* is the ratio between the standard deviations for a particular micro-cluster *i*. We then report the distribution of ratios across all micro-clusters for each protein separately, as presented in Supplementary Figure 1b.

### Cell-Cell Similarities from Trajectories

VISION interfaces with the Dynverse package [12] to process trajectories in an analysis pipeline. The results of running a trajectory with Dynverse is an abstracted trajectory model which VISION is able to ingest and process. Most essential to the VISION pipeline are two components of the Dynverse model: (a) the “milestone” network detailing the topology of the trajectory (e.g., in a developmental process, milestones would be important cell states or types and the topology would represent how these states are related to one another) and (b) the progress of cells along this network (i.e., where cells lie between the important milestones).

Using the milestone network and the progress of cells between each pair of milestones (i.e. a “pseudotime”) we define cell-cell similarities according to the tree-based geodesic distances. Given this cell-cell similarity map, we can then perform the same autocorrelation score evaluation for all signatures as described above.

VISION visualizes the trajectory by first applying a method to visualize the milestone network and then projecting the cells onto their assigned edges, where their locations between edges are proportional to their pseudotime. VISION uses a variety of methods for visualizing the milestone network such as Fructerman-Reingold and Davidson-Harel. Importantly, to help visualize edges where many cells are located, we add a small amount of jitter to each cell’s position perpendicular to its assigned edge.

### Differential Signature Analysis

Similar to a differential gene expression test, VISION performs a test to identify which signatures’ scores are differential among a particular group of cells. These groups of cells are defined using any input meta-data of a categorical nature (i.e. discrete variables such as disease status or clustering assignments). For each supplied categorization, we test for signatures that are differential, by performing a Wilcoxon rank-sum test for every 1 vs. All comparison. The results of these tests represent one of the “label-based” analyses performed by VISION and are available for browsing in the output VISION report.

### Autocorrelation Score of Discrete Meta-Data

The Geary’s C cannot be used to evaluate the autocorrelation of discrete meta-data variables (such as donor or batch), and so instead, VISION uses a procedure based on the chi-squared test. First the local distribution of the variable is computed around each cell. Then, these local distributions are aggregated into a square contingency whose rows represent the distribution of the variable as observed local to the cells’ of each value. For example, if run on a batch variable, the row representing batch *x* will contain proportions of each batch as estimated from the local neighborhoods of cells in batch *x*. This table is then evaluated with the chi-square test.

More concretely, first, for each cell, *i*, a local proportion for each variable value *m* is evaluated as:

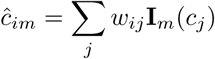

Here, the weights *w_ij_* are computed from the manifold using the same procedure described above for transcriptional signatures, *c_j_* represents the value of the discrete variable of interest, and I*_m_*(*x*) is an indicator function that takes on a value of 1 if *x* = *m* and 0 otherwise. From these values, the contingency table X is computed as:

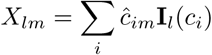

The chi-squared test is then computed on this contingency table X to estimate a p-value.

Because of the large number of cells involved in modern scRNA-seq experiments, it is possible to achieve a significant p-value for an autocorrelation effect that is too weak to be of interest. Accordingly, VISION also reports the effect size as the Cramer’s V in addition to the p-value. This indication of effect size ranges from 0 (no autocorrelation) to 1 (perfect autocorrelation), and provides an additional means to rank and categorize potentially confounding effects.

### Analysis of Single-Cell Expression Profiles from Lupus Cohort

Peripheral blood mononuclear cells were isolated from 16 patients (12 diagnosed with SLE and 4 healthy controls), Ficoll separated, and cryopreserved by the UCSF Core Immunologic Laboratory (CIL). Cellular suspensions were loaded onto the 10x Chromium instrument (10x Genomics) and sequenced as described in Zheng et al. [3]. This collection represents a single batch (batch “7.19”) of a larger study (manuscript in preparation).

Counts per gene were quantified using the CellRanger pipeline by 10x Genomics and the hg19 human genome reference. For cases where multiple Ensembl IDs had the same gene symbol, counts were combined by summation. Genes with less than 10 counts were removed from downstream analysis. Scaled counts were produced by dividing the counts in each cell by the total number of UMI detected for that cell and multiplied by 4000 (approximately the median number of UMI’s per cell). Data was further normalized by linearly regressing out the Number of UMI’s per cell from the log-transformed scaled counts. We used the CellRanger output .bam file directly as input to the algorithm demuxlet with default settings to identify the donor of origin for each cell [47].

To select the B cell cluster used in this analysis, we first clustered the cells in an unsupervised manner and then used cell-type signatures in VISION to identify the cluster corresponding to B cells (Supplementary Figure 2a,b). To cluster the cells, the normalized expression matrix was first transformed via PCA (20 components), and then clustered using the louvain algorithm. The subset of the normalized expression matrix corresponding to this B cell cluster was then used as input into VISION.

To further test the association between the SLEDAI score and the Interferon-Alpha signature score we subset the B-cell cluster (shown in Supplementary Figure 2A) selecting only the cells from SLE donors. B cell signature scores were averaged within each donor and these donor-averages were tested against donor SLEDAI scores using a one-sided Kendall tau correlation test against an empirical background of shuffled values (100k replicates). The same method was used to test the TNF-Alpha signature values yielding an insignificant result.

### Analysis of HSCs

The expression profiles of 5,432 Hematopoietic Stem Cells (HSCs) were obtained from NCBI GEO, accession GSE89754; in this analysis, we used the raw UMI counts of the basal bone marrow HSCs (specifically, GSM2388072) [39]. To compute the trajectory, we first filtered the genes using the gene set that the original authors used, and removed cells which the authors flagged as not passing their own internal filters. Monocle2 [7] was used to compute the trajectory, where we used the “log” normalization scheme, and wrapped the final inferred trajectory with dynverse [12]. The cell types reported here are those used in the original study.

We then scaled the raw UMI counts such that each for the expression level of gene *g* in cell *c* equaled

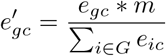

where we have *G* genes and *m* is the median number of counts that are observed across all cells in the dataset. With this transformation, we have scaled each cell such that it has a total of *m* UMIs.

A VISION object was created with these scaled counts, signatures consisting of both the Hallmark and C2 MSigDB collections [20], and the Dynverse-wrapped Monocle trajectory.

### Comparison of VISION to Existing Tools

To broadly convey the unique features of VISION, we conducted a qualitative comparison of VISION to other similar tools that seek to combinations of functional analysis and visualization for large scRNA-seq datasets. Supplementary Table 1 summarizes this comparison for a panel of methods, including Spring ([43]), CCS ([48]), ROMA ([46]), PAGODA, ([45]), MAST ([44]), Scanpy ([42]), and Seurat ([41]). Evidently, VISION has a comprehensive set of analysis capabilities, some of which (e.g., annotating trajectories or adding meta data to the analysis) are unique whereas others (e.g., performing cluster-based, but not cluster-free analysis) are only partially present in other packages.

A key distinguishing feature of VISION is its ability to annotate the biological meaning of cell to cell variability both with and without the need to first partition cells into groups. While most existing tools are restricted to the former type of analysis, it is important to note that with single-cell datasets, often cells do not neatly partition into groups, and instead the variation within a group is of primary interest (see example in Figure 2). Out of the reference tools surveyed above, two methods – ROMA ([46]) and PAGODA ([45]) were designed to identify and annotate important axes of biological variation in a dataset without the need for a-priori stratification of the cells. To demonstrate the value of VISION compared with these methods, we ran both ROMA ([46]) and PAGODA ([45]) on the B-cell cluster presented in our manuscript with the Hallmark (MSigDB [20]) signature set. ROMA did not select any of the signatures as significant despite the clear coordinated variation exhibited in the TNF*α* and INF*α* response pathways. To further evaluate VISION vs. PAGODA, we compared the performance of the two methods on expected sources of biological variation in this sample. The first source of variation is that these B cells are from a mix of Lupus-diagnosed (SLE) and healthy individuals. One signature, from the MSIGDB c7 collection, is specifically designed to capture this contrast (based on SLE studies [49] and [50], see GEO entry GSE10325). Supp. Figure 5a shows that the signature scores from VISION better reflect this expected dichotomy by showing a larger distributional difference between healthy and SLE patients than those of PAGODA (ROC AUC 0.79 vs 0.61). As this is a major source of variation in the data, we would expect other signatures to distinguish these two donor groups. By taking a diverse collection of signatures from MSIGDB (Hallmark, C2 Canonical Pathways, and C7 restricted to B Cell signatures) and NetPath and repeating the analysis above (computing a ROC AUC for a classifier based on each signature’s scores), it is observed that the signature scores of VISION tend to better differentiate cells from the healthy cohort vs. the SLE cohort (Supp. Figure 5b). Another expected axis of biological variation in this B cell sample is the distinction between naive and memory subtypes, which can be discerned by looking at marker genes (see Supp. Figure 6a). By visualizing naive vs. memory B cell gene signature scores (e.g., based on [51]; GEO entry GSE11386) from both PAGODA and VISION it is evident that VISION signature scores are more likely to correctly identify this variation and VISION additionally selects these signature scores as being statistically significant (Supp. Figure 6b).

**Supplementary Figure 1:**
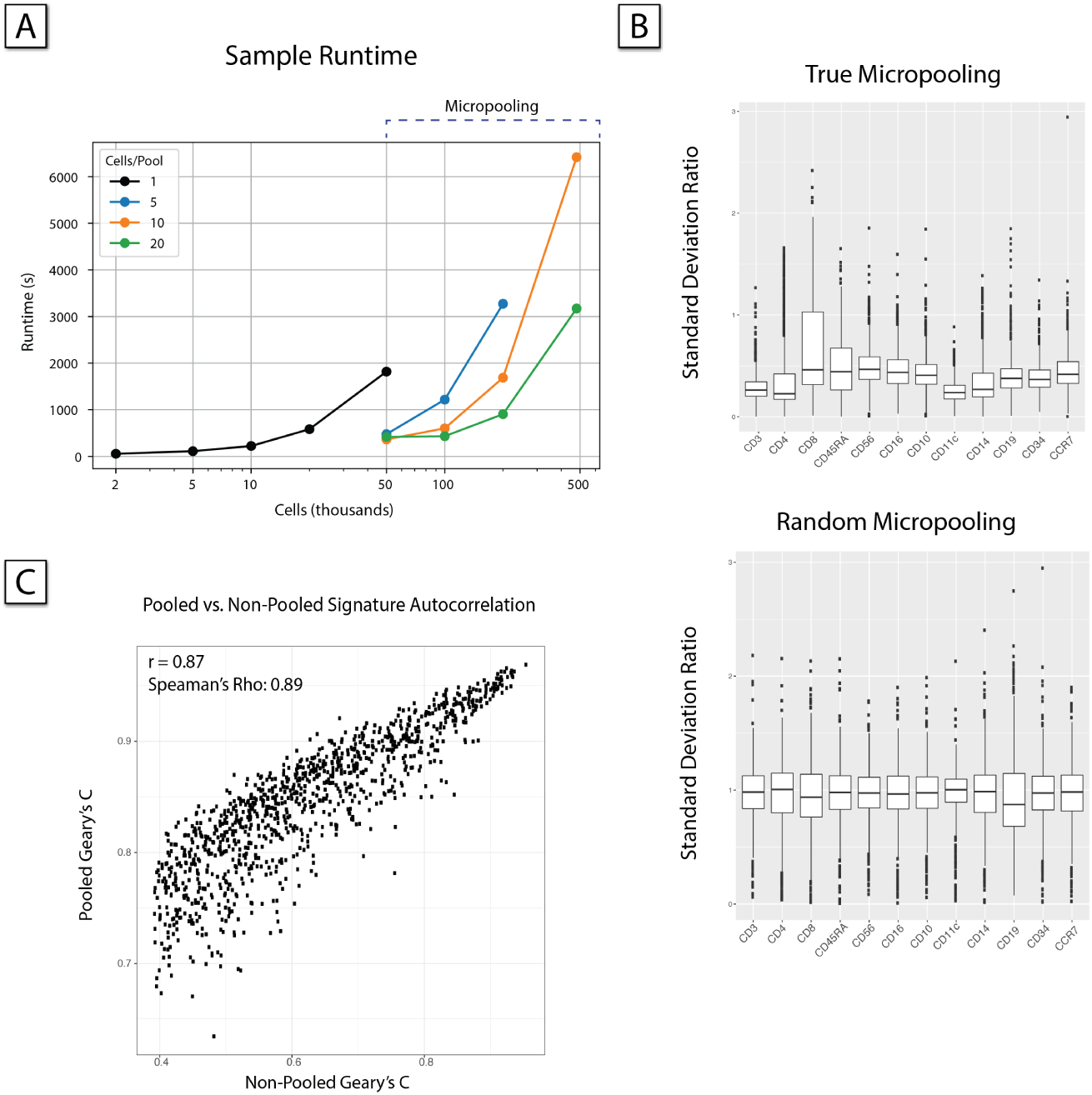
Micro-pooling allows VISION analysis to comfortably scale to very large cell counts. A) Sample runtime for the full pipeline (10 cores). B) To confirm that similar cells were being clustered, we used 9,000 Cord Blood Mononuclear Cells (CBMCs) whose mRNA and protein abundances were profiled simultaneously with the Cite-seq protocol [23]. We find that the cells in each micro-cluster are biologically coherent - specifically, the variation in measured surface protein abundance within micro-clusters is much less than the variation within randomly partitioned micro-clusters. C) The signature autocorrelation scores are compared before and after micro-pooling.

**Supplementary Figure 2:**
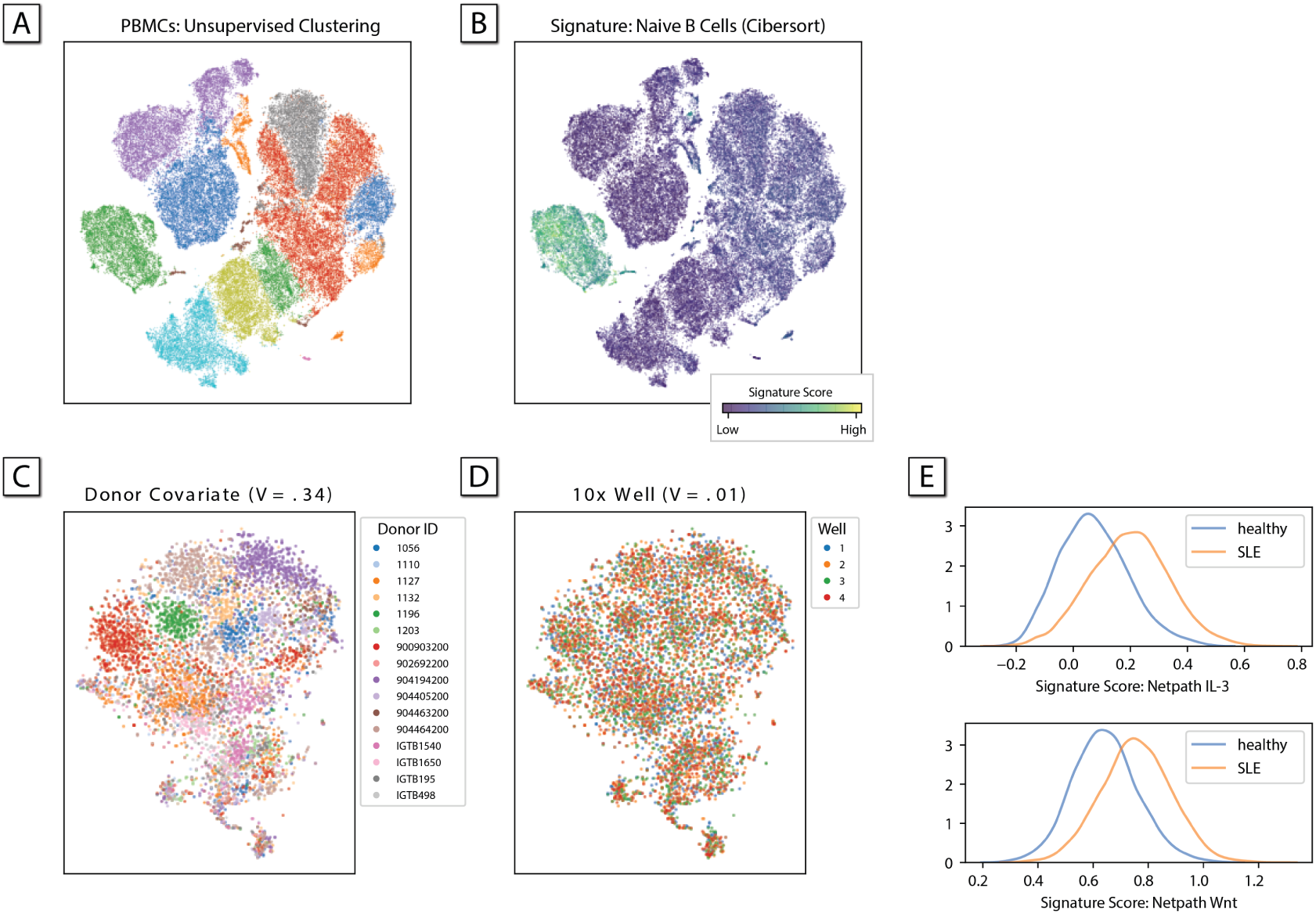
A) 67,000 PBMCs from a mixed (healthy/SLE) cohort of 16 individuals. Cells have been placed into unsupervised clusters by VISION via the louvain algorithm. B) Alternately, label-based (differential) analysis highlights signatures which help annotate clusters. Shown here, the Naive B cell signature from CiberSort [52]. C) B cell cluster from the cell’s in panel (A). VISION quantifies the degree to which covariates (shown: donor) contribute to cellular heterogeneity. D) Well covariate is visualized in the same set of B Cells. The effect size is small (V=0.01) in contrast to the effect size of the donor covariate in panel (D). E) Additionally, VISION quantifies the degree to which signatures stratify according to covariate label. Shown here, top signatures derived from the NetPath library whose scores discriminate healthy from SLE patients implicate the IL-3 and Wnt signaling pathways.

**Supplementary Figure 3:**
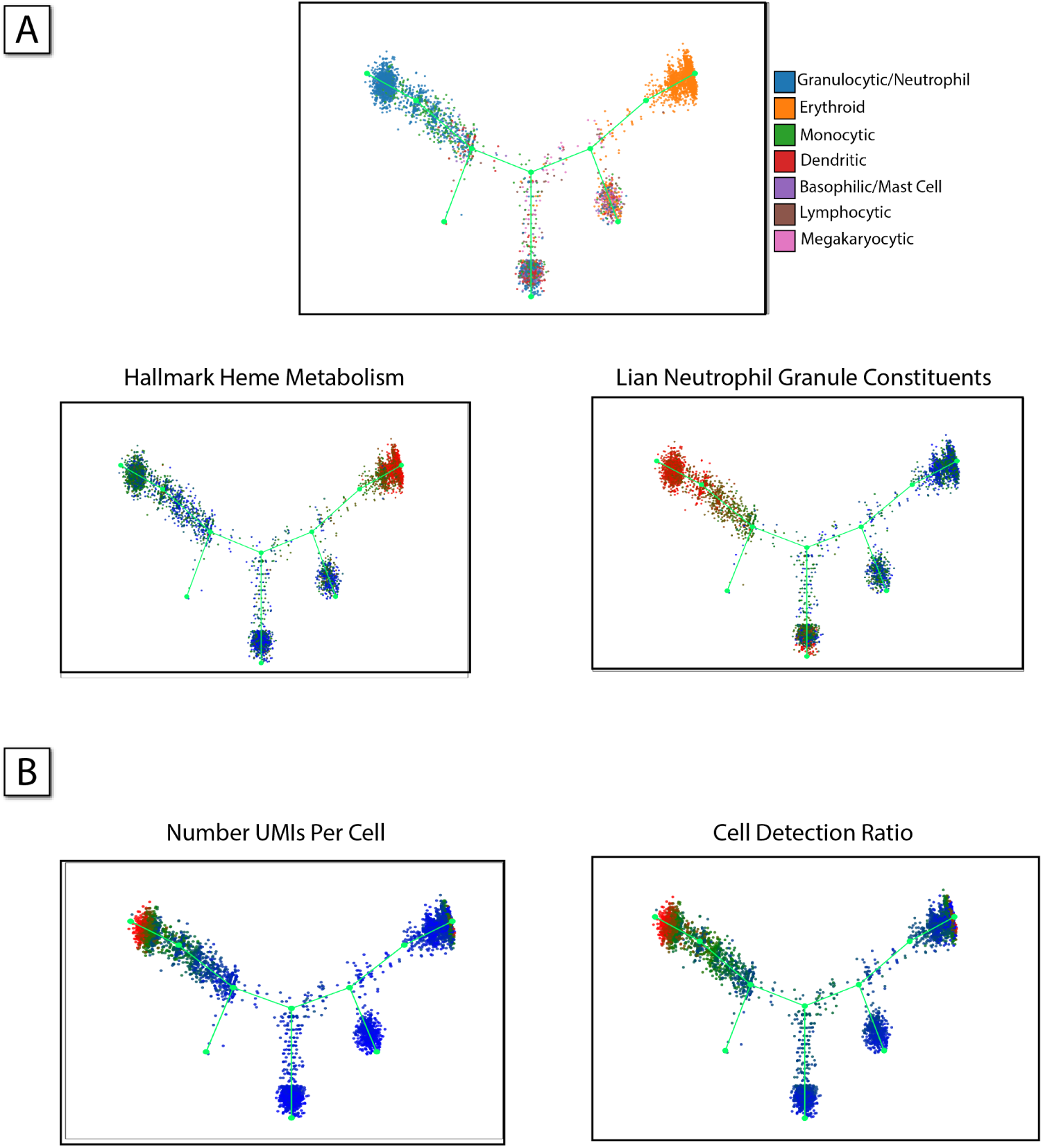
Cell-type specific signatures differentiate branches of an inferred haematopoietic trajectory. (A) Trajectory representation of cells undergoing hematopoiesis, colored by the cell type’s inferred by Tusi et al. [39] in the original study. Hallmark Heme Metabolism, a cell-type signature for the erythrocytic lineage, aligns well with where erythrocytes are found in the trajectory. Lian Neutrophil Granule Constituents, a cell-type signature for the granulocytic lineage, aligns well with where granulocytes are found in trajectory. (B) Cell level meta-data can be used as a data-driven signature; in this case, the number of UMIs and cell detection ratio (CDR; the ratio of detectable genes per cell) are strikingly localized to the granulocyte arm likely due to their increased diameter (16 *μ*m vs ~ 8*μ*m) compared to other white blood cells and erythrocytes.

**Supplementary Figure 4:**
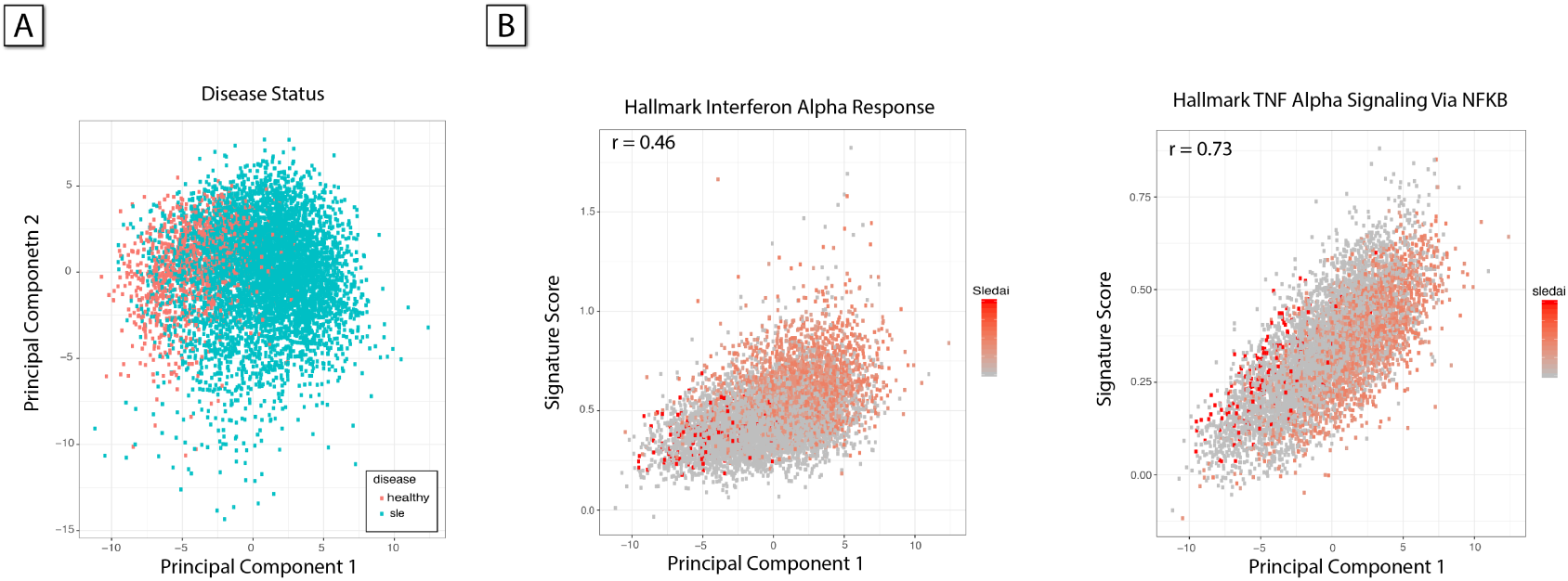
The PCAnnotator highlights sources of biological heterogeneity by calculating the correlation between signatures and expression principal components. (A) Disease status strikingly is stratified by the first principal component, indicating that signatures well correlated with the first principal component may be sources of this stratification. (B) Two signatures with high local autocorrelation scores are highly correlated with the first principal component - Hallmark Interferon Alpha Response (*ρ* = 0.46, *p* < 10^−3^) and Hallmark TNF Alpha Signaling Via NFKB (*ρ* = 0.73, *p* < 10^−3^).

**Supplementary Figure 5:**
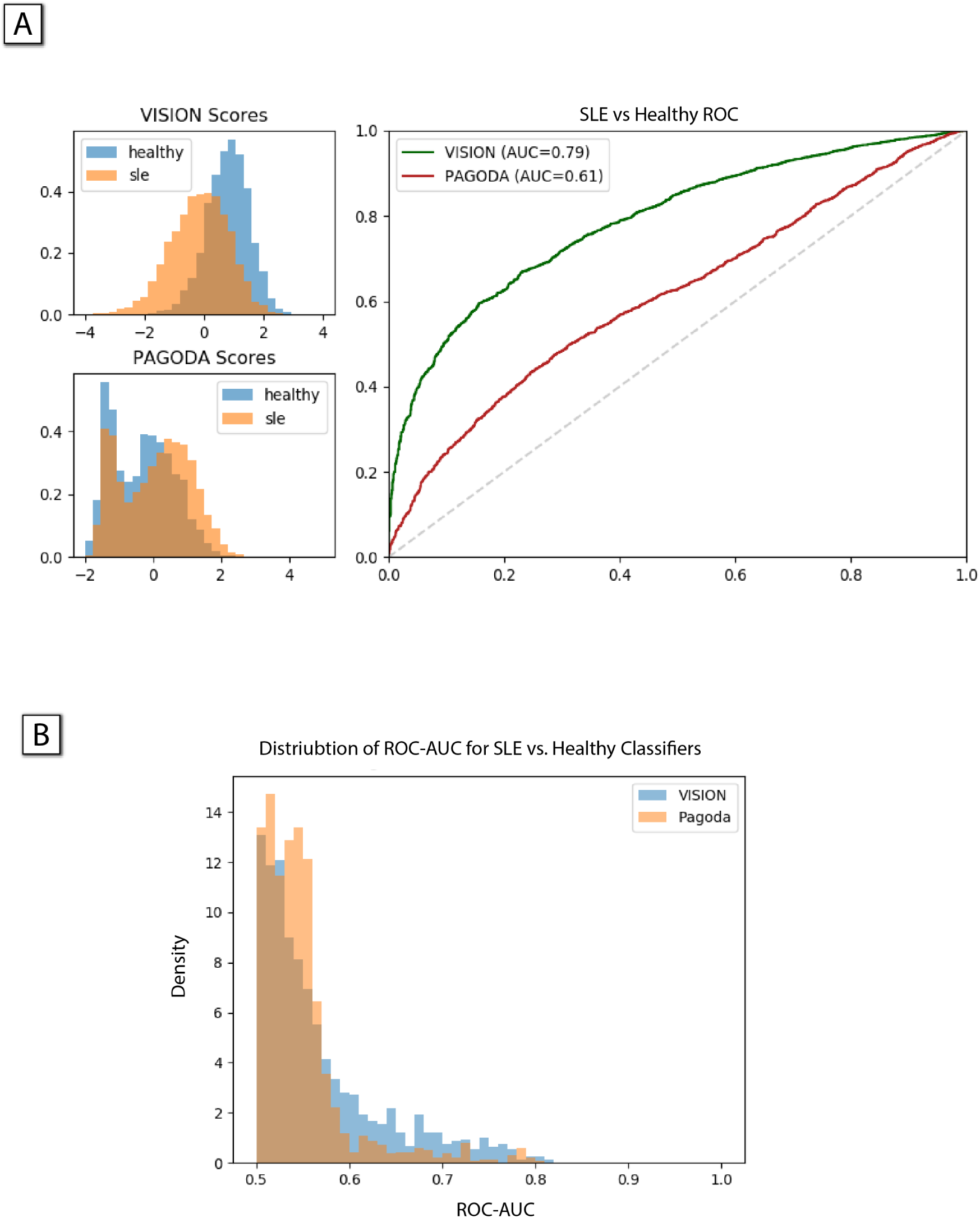
VISION’s signature scores more reliably capture underlying biology.(A) Using a signature from the MSIGDB c7 collection, GSE10325_BCELL_VS_LUPUS_BCELL, the disease status can be predicted. Signature scores for this signature from PAGODA and VISION were used to compare the effectiveness of these scores at capturing biological signal; we find that VISION has an AUC-ROC of 0.79 and PAGODA has an AUC-ROC of 0.61, indicating that VISION is more effective in this respect. (B) The distribution of AUC-ROC’s when classifying disease status of single cells for a diverse set of signatures illustrates that VISION overall is more capable of differentiating cells from the healthy cohort vs the SLE cohort.

**Supplementary Figure 6:**
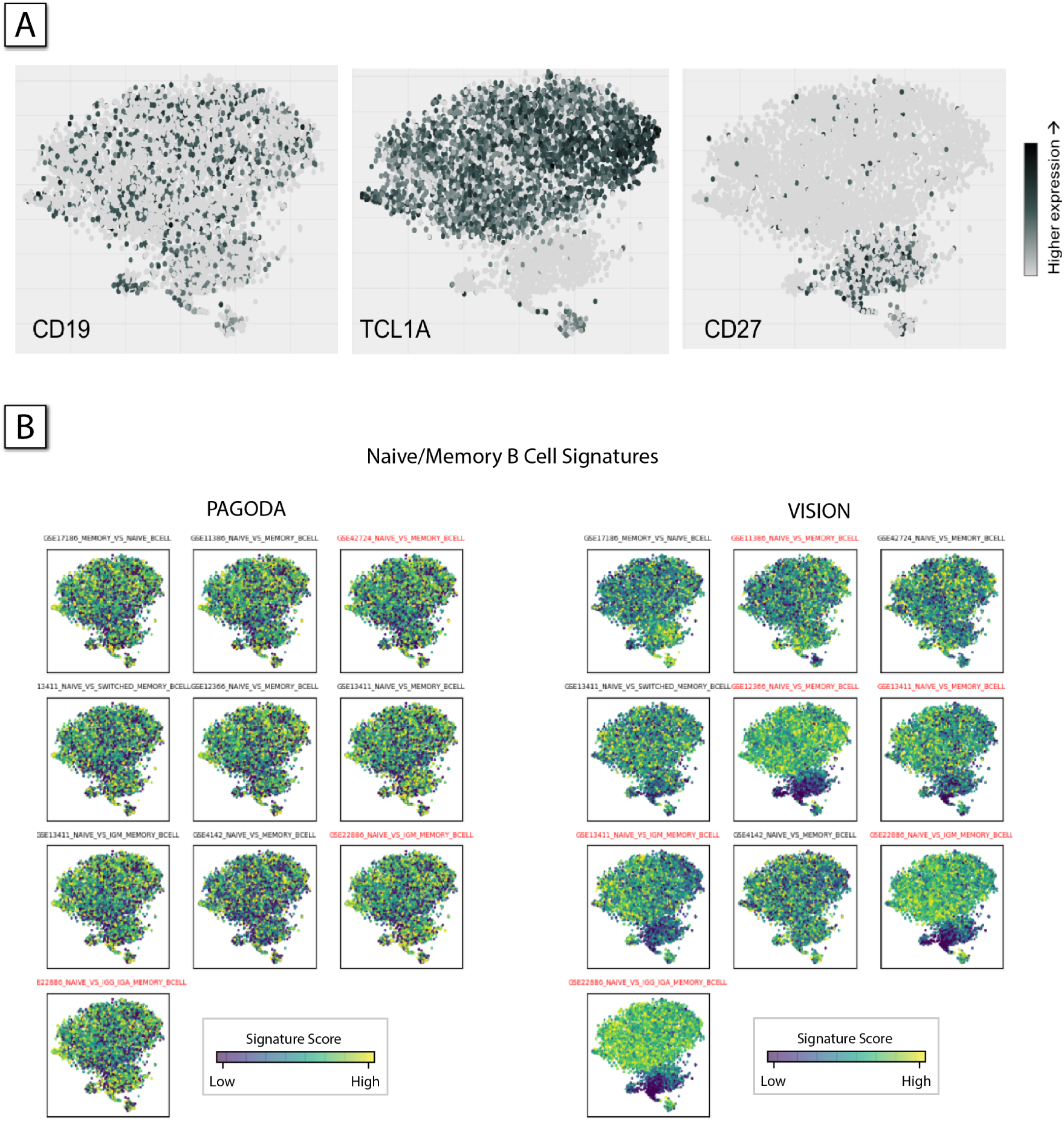
(A) Expression of individual genes in B cells from a cohort of healthy and SLE donors. Evidently, the expression of CD19 is prevalent and present throughput the B cell population. Conversely, the expression of CD27, a canonical marker for memory B cells, is restricted to a sub-cluster, interpreted by VISION to be a subpopulation of memory B cells (see B). Similarly, the expression of TCL1A, a marker for naive B cells is not expressed in that cluster. (B) Signatures which describe the naïve vs. memory dichotomy in B cells are visualized. Signatures marked as significant PAGODA (left three columns) or VISION (right three columns) are titled in red. While there is some overlap, the signatures denoted by VISION more consistently distinguish the two main clusters. Highly ranked (significant) signatures are marked in red font.

**Supplementary Table 1:** VISION has several important properties that distinguish it from other software packages for automated annotation and for visualization and exploration of single cell data. First, VISION has a comprehensive set of data analysis capabilities. Some of these capabilities (e.g., annotating trajectories or adding meta data to the analysis) are unique to VISION; other properties are only partially present in other packages (e.g., performing cluster-based, but not cluster-free analysis).

|  | Spring | CCS | Roma | Pagoda | MAST | Scanpy | Seurat | Vision |  |
| --- | --- | --- | --- | --- | --- | --- | --- | --- | --- |
| Evaluation of Pathway Scores |  |  | ✓ | ✓ |  | ✓ | ✓ | ✓ | Analysis |
| Differential Pathway Analysis |  | ✓ |  |  | ✓ |  |  | ✓ |  |
| Confounding of Factor Meta-Data |  |  |  |  |  |  |  | ✓ |  |
| Confounding of Numerical Meta-Data |  |  |  |  |  |  |  | ✓ |  |
| Label-free signature rankings |  |  | ✓ | ✓ |  |  |  | ✓ |  |
| Allows input of Latent Space |  |  |  |  |  |  |  | ✓ |  |
| Allows input of Trajectories |  |  |  |  |  |  |  | ✓ |  |
| Visualize Latent Space |  |  |  |  |  | ✓ | ✓ | ✓ | Visualization |
| Visualize Trajectory | ✓ |  |  |  |  | ✓ |  | ✓ |  |
| Dynamic Web Visualization | ✓ |  |  | ✓ |  |  |  | ✓ |  |

